# C Terminus of DJ-1 Determines Its Homodimerization, Deglycation Activity and Suppression of Ferroptosis

**DOI:** 10.1101/2020.03.26.992446

**Authors:** Li Jiang, Xiaobing Chen, Qian Wu, Haiying Zhu, Chengyong Du, Meidan Ying, Qiaojun He, Hong Zhu, Bo Yang, Ji Cao

## Abstract

DJ-1 is a multi-functional protein related to cancer and autosomal early-onset Parkinson disease (PD). Besides the well-documented antioxidative stress activity, recent studies suggest that DJ-1 has the deglycation enzymatic activity and the anti-ferroptosis function. Although it has been demonstrated that DJ-1 forms the homodimerization, which dictates its antioxidative stress activity, the relationship between the dimeric structure and newly reported activities remains largely elusive. In this study, we find that the deletion mutation of the last 3 amino acids at C terminus of DJ-1 disrupts its homodimerization in both transfected and purified DJ-1 protein. Further study shows that hydrophobic L187 residue is of great importance for DJ-1 homodimerization. In addition, the ability in methylglyoxal detoxification is almost abolished in the mutation of deleting last 3 residues at C terminus (ΔC3) and point mutant L187E compared with wild type DJ-1 (DJ-1 WT). We also find that the suppression of ferroptosis is fully inhibited by ΔC3 and L187E while partially suppressed by V51C. Thus, our findings show that C terminus of DJ-1 is crucial for its homodimerization, deglycation activity and suppression of ferroptosis.

## Introduction

DJ-1 is a highly conserved multi-functional protein first identified as an oncogene synergetically transforming mouse NIH3T3 cells with activated H-Ras and C-Myc [1]. It is localized predominantly in cytoplasm and partially in mitochondria, and it also expresses in the nucleus of several cell types [2]. It has been reported that DJ-1 is over-expressed in multiple tumor cells and is positively related to tumor metastasis and poor prognosis in patients, which further suggest it’s a potential predictive biomarker for cancer diagnosis and prognosis [3–5]. Moreover, knockdown of DJ-1 promotes the efficacy of chemotherapeutic drug in some tumor cells [6]. The regulatory mechanisms of oncogenic transformation by DJ-1 is reported to associated with inactivation of phosphatase and tensin homolog (PTEN) and activation of c-Raf [7, 8]. Besides, the well-known effects of DJ-1 on antioxidative stress might be essential for tumourigenesis and cancer progression [6, 9]. DJ-1 undergoes irreversible oxidation at the Cys106 residue to directly eliminates reactive oxygen species (ROS) [10–12]. Mechanistically, the role of DJ-1 as an abundant antioxidant scavenger of soluble ROS (such as H2O2) is related to the antioxidant transcriptional master regulator nuclear factor erythroid 2-related factor 2 (NRF2), which orchestrates the expression of genes coding for stress response and antioxidant proteins [9, 13]. In addition, we have shown that DJ-1 function as a key factor protecting tumor cells by affecting both soluble reactive oxygen species (ROS) and lipid ROS with distinct mechanisms [14, 15]. The different oxidation states of DJ-1 determines the cell fate via either activating autophagy or apoptosis through regulation of the activity of ASK1, which is based on the activity of DJ-1 against soluble ROS [15]. While the negative effects of DJ-1 on ferroptosis is not attributed to its classical role in stabilizing NRF2. DJ-1 suppresses ferroptosis triggered by lipid ROS through maintaining cysteine synthesized from the transsulfuration pathway, the predominant source of cysteine for GSH biosynthesis when the cystine/glutamate transporter (SLC7A11, xCT) is inhibited [14]. Moreover, the suppression of the DJ-1 expression dramatically enhanced the sensitivity to the ferroptosis inducer Erastin of tumour cells both *in vitro* and *in vivo* [14]. Thus, DJ-1 is further proposed as a potential therapeutic target for cancer therapy.

Though DJ-1 is initially reported as an anti-oxidative enzyme through directly reducing mixed disulfides in covalently modified proteins, several clues of evidence also suggest that DJ-1 has the chaperon activity and can be converted from a zymogen to a protease by carboxyl-terminal cleavage [16, 17]. Interestingly, most recent studies showed that DJ-1 has glyoxalase enzymatic activity *in vitro*, converting glyoxal (GO) or methylglyoxal (MGO) into glycolic or lactate, respectively [18]. And later, it was recognized as an apparent activity that in fact reflects its deglycase activity catalyzing the deglycation of the Maillard adducts formed between amino groups of proteins or nucleotides and reactive carbonyl groups of glyoxals [19, 20]. Thus, cells depleting DJ-1 display increased levels of glycated DNA, DNA strand breaks, and phosphorylated p53, which provides another layer to understand the promoting effect of DJ-1 knockdown on the efficacy of chemotherapeutic drug in some tumor cells [21]. Therefore, deglycation activity of DJ-1 might provide another valuable direction for the function related to cytoprotective effects of DJ-1 and the compounds inhibiting the deglycation activity of DJ-1 might offer new opportunities to facilitate cancer therapy [10, 22].

The crystal structure of DJ-1 shows that it exists as a dimer comprising 11 β-sheets (β1-β11 sheet) and 8 α-helices (αA-αH helix) [23, 24]. The helix-sheet-helix motif presents a sandwiched structure [24]. The αA-helix motif is located at the center of the dimer structure, while αH-helix motif is at the C terminus of DJ-1 [23–25]. It is reported that the C-terminal G-helix-kink-H-helix motif is essential for protein stability and survival promoting activity of DJ-1 [26]. The L166P point mutant in G-helix motif of DJ-1 is one of the most striking mutations which impairs dimer formation and dramatically accelerates DJ-1 degradation [27–29]. Although more attention has been given to the C-terminal G-helix-kink-H-helix motif of DJ-1, the more accurately relationship between its C terminus, especially the last three amino acids and its homodimerization and biological functions remains unclear.

The aim of this study is to specifically investigate the structure basis of dimeric DJ-1 and its role in deglycation activity and ferroptosis. We found that C terminus of DJ-1 is crucial for its dimeric formation and biological function, regulating its homodimerization, deglycation activity and suppression of ferroptosis. The results of this study provided an important insight that will fuel the future development of DJ-1 inhibitor targeting homodimerization as promising therapeutics for cancer therapy.

## Materials and Methods

### Antibodies and Reagents

The antibody to DJ-1 (#5933) was obtained from Cell Signaling Technology (Boston, America). The primary antibodies against GAPDH (db106), Myc (db2603) and Flag (db7002) were obtained from Diagnostic Biosystems (Hangzhou, China). MGO (W296902), GSH (G4251), NAC (A7250) were obtained from Sigma-Aldrich (St. Louis, America). Erastin (S7242) was obtained from Selleck Chemicals (Houston, America). DMEM (GIBICO, #12800) was obtained from Thermo Fisher Scientific (Waltham Mass, America).

### Cell Culture

The HEK293T cell line was purchased from the Shanghai Institute of Biochemistry and Cell Biology (Shanghai, China). The MEF DJ-1 KO cells were produced from day 13.5 embryos of DJ-1 KO mice (B6.Cg-Park7tm1Shn, #006577, The Jackson Labratory) according to standard procedures. HEK293T and MEF DJ-1 KO cells were cultured in DMEM medium. All mediums contain 10% fetal bovine serum (Hyclone, SV30160.03, GE Healthcare), 100 units per mL penicillin and 100 μg/mL streptomycin. Cells were incubated at 37°C in a humidified atmosphere of 5% CO2 and monitored for mycoplasma contamination every six months. HEK293T cells were authenticated by STR profiling.

### DJ-1 Expression and Purification

The cDNA of DJ-1 and their mutants were cloned into pET28a vectors, which express N-terminal 6×His-tagged DJ-1 (the primers are listed in table S1). The cloned vectors were transformed into E.coli *Trans BL21* (DE3) (CD601-02, TransGen Biotech), which were grown at 37°C in LB broth containing 0.1 mg/ml of Kanamycin until the OD600 reached 0.4-0.6. Isopropyl-b-D-thiogalactoside (IPTG, 0.25 mM) (V900917, Sigma-Aldrich) was then added to over-produce protein, and bacteria were further incubated at 25°C for 16 h. After centrifugation, bacteria were resuspended with lysis buffer (20 mM Na3PO4, 500 mM NaCl, 50 mM imidazole, pH 7.4), homogenized by high pressure cell cracker and centrifugated (8,000g) at 4°C for 30 min. The supernatant was loaded onto the Ni^2+^-NTA column and the protein was eluted with 500 mM imidazole. The purified proteins were dialyzed in 100 mM Na3PO4 (pH 8.0) buffer containing 20% glycerinum.

### Lentivirus Transduction

The pCDH-EF1-Puro plasmid was obtained from System Biosciences. The pCDH-DJ-1 WT and mutants were constructed using the primers listed in table S2. Lentivirus was generated by HEK293T cells with pCMV-R8.91 (packaging vector), pMD2-VSVG (envelope vector) and pCDH plasmids co-transfection using Lipofectamine 2000 (#11668019, Invitrogen). Medium containing virus was harvested 48 h after transfection and filtered by 0.45 μm millipore filter. Cells were grown in 6-well plates at 20% to 30% confluency for infection, and 0.1-1 mL of each virus was added with 2 μL polybrene (6 mg/mL). After infection for 16 h, the medium was changed to fresh medium.

### Western Blot Analysis

Cells were lysed with 1% NP40 buffer (50 mM Tris-HCl, 150 mM NaCl, 1% NP40, pH 7.4, 0.1 mM sodium vanadate, 5 μg/mL leupeptin, 0.1 mM phenyl methane sulfonyl fluoride) and incubated at 4°C for 30 min, flicked every 10 min to fully lysed. The lysate was then centrifuged at 12,000 g for 30 min at 4°C to remove insoluble materials. Protein concentrations of wholecell lysates were determined by the QuantiPro™ BCA Assay Kit (Sigma-Aldrich, QPBCA). The proteins were then electrophoresed in SDS-PAGE gels to separate by molecular weight and transferred to polyvinylidene difluoride membranes (Millipore, Bedford, MA, USA). After blocked with 5% non-fat milk, the membrane was incubated with suited primary antibody overnight at 4°C and followed by HRP-labeled secondary antibodies. And proteins were visualized using enhanced chemiluminescence detection (NEL103E001EA, PerkinElmer) by AI600 (GE Healthcare).

### Immunoprecipitation

Exogenous immunoprecipitation was performed by anti-Flag IP resin (L00425, GenScript) or anti-Myc magnetic beads (B26201, Bimake). Briefly, after determined by the QuantiPro™ BCA Assay Kit (QPBCA, Sigma-Aldrich), 500 μL whole-cell lysates were added into the matched beads for 8 h at 4°C with end-over-end mixing. Finally, the complex was washed with T-PBS for five times and then suspended in loading buffer and followed by western blot analysis.

### DSS-Mediated Cross-Linking Assay

Whole-cell lysates were determined and incubated at 4°C for 30 min in 2 mM DSS crosslinker (Suberic acid bis (N-hydroxysuccinimide ester), #S1885, Sigma), using isometric DMSO as control. The reaction was quenched with 1 M Tris-HCl buffer (pH 7.4) for 15 min at room temperature. Samples were analyzed by western blot.

### Size Exclusion Chromatography (SEC)

100 μL of purified recombinant protein (at approximately 2 mg/mL) were loaded onto a size-exclusion chromatography column (PL1580-3301, Agilent) in the phosphate buffer (20 mM Tris, pH 8.0, 150 mM NaCl, pH 7.4) at 0.35 mL/min with an high performance liquid chromatograph (1260 Infinity II, Agilent).

### Glyoxalase Assay

The purified wild type DJ-1 protein or its mutants (6 μM) were mixed with MGO (6 μM) in 50 mM sodium phosphate buffer (pH 7.0) at a total reaction volume of 70 μl at room temperature. The reaction was stopped by adding 120 μl 0.1% 2,4-dinitrophenylhydrazine (DNPH) solution. The solution was incubated for 15 min at room temperature. Then 160 μl 10% NaOH was added. After further incubation for 15 min, absorbance (540 nm for MGO) was measured.

### Deglycase Assay

Experiments were performed in 50 mM sodium phosphate buffer (pH 7.0) at room temperature. MGO and N-acetylcysteine were premixed to a final concentration of 20 mM. After reaching steady-state absorption levels at 288 nm, recombinant DJ-1was added to a final concentration of 2 μM. An identical volume of sodium phosphate buffer was added as controls. Absorbance at 288 nm for hemithioacetal was measured for 60 min.

### Analysis of Lipid ROS Production

Cells were harvested by trypsinization, resuspended in 500 μL PBS containing 2 μM C11-BODIPY (581/591) (#D3861, Invitrogen) and incubated for 30 min at 37°C in an incubator. Cells were then resuspended in 500 μL fresh PBS, filtered through a 40 μm cell strainer, and analyzed using a flow cytometer (FACSuite, BD Biosciences) equipped with a 488 nm laser for excitation. Data were collected from the FL1 channel (527 nm). For each sample, at least 10,000 cells were analyzed.

### Cell Viability Assay

Cell viability was typically assessed in 96-well format by Cell Counting Kit-8 (CCK8) (HY- K0301, MedChem Express). When added to cells, WST-8 [2-(2-methoxy-4-nitrophenyl)-3-(4-nitrophenyl)-5-(2, 4-disulfophenyl)] is modified by the reductive product of viable cells and turns orange (WST-8 formazan) in color. Briefly, 10 μL of the CCK-8 solution was added to each well of the plate, incubated for 1-4 h at 37°C, and the absorbance at 450 nm was measured on a microplate reader (Multiskan spectrum 1500, Thermo). Cell viability of test conditions was reported as a percentage relative to the negative control.

### Statistical analysis

All statistical analyses were performed using Prism 7.0c (GraphPad Software). The number of biological replicates for each experiment is indicated in the figure legends. Values are presented as means ± standard deviation (SD). Differences between means were determined using the two-tailed, unpaired *Student’s t tests*, and were considered significant at p < 0.05.

## Results

### The C terminus determines dimeric DJ-1 formation

To investigate the dimerization of DJ-1, we first established stable DJ-1 over-expressed HEK293T cells using Flag-tagged and Myc-tagged DJ-1 lentivirus. The ectopic DJ-1 was purified from cells by immunoprecipitation. Our data illustrated that Myc-tagged DJ-1 could be pulled down by Flag-tagged DJ-1 (Fig. 1a) and the same result also was observed by pulling down the Myc-tagged DJ-1 (Fig. 1b), indicating that DJ-1 has dimerization. Then we studied the homodimerization of DJ-1 by disuccinmidyl suberate (DSS), an intramolecular crosslinking agent which could make polymerized protein avoid disassembly in reductant such as β-mercaptoethanol during the western blot analysis. When cell lysate was treated with DSS for 30 min at 4°C, we found that most protein represented as dimer both in Flag-tagged or Myc-tagged DJ-1 (Fig. 1c). In addition, the dimerization was also detected in Flag-tagged and Myc-tagged DJ-1 co-expression system, suggesting that the N terminus tag doesn’t affect the dimeric DJ-1 formation (Fig. 1c).

**Fig. 1.**
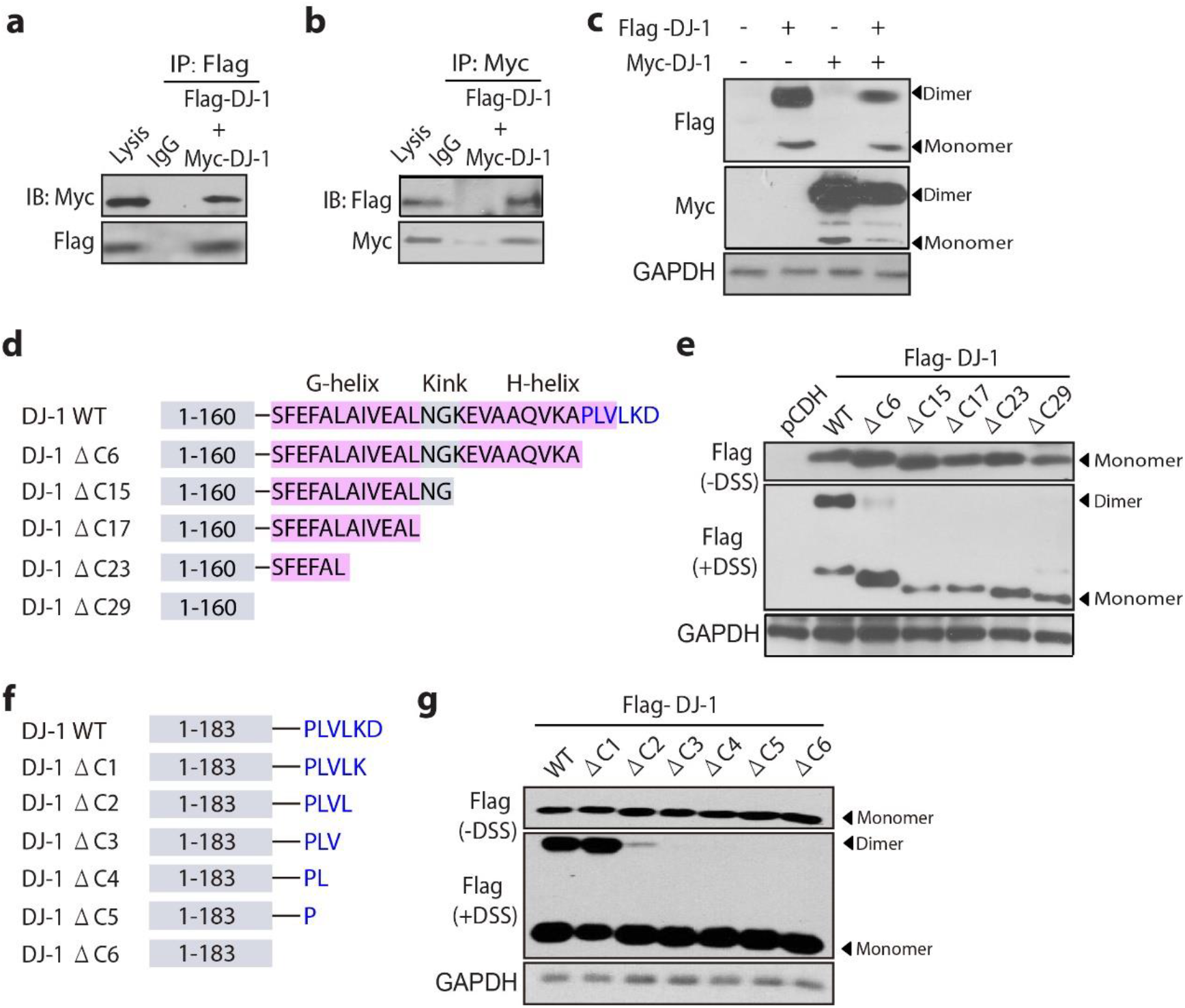
The C terminus determines dimeric DJ-1 formation. (**a-b**) The interaction between Flag-tagged and Myc-tagged DJ-1 was analyzed to represent the dimerization of DJ-1 protein. Indicated HEK293T cells stably expressing Flag-tagged and Myc-tagged DJ-1 were harvested for immunoprecipitation and subjected to immunoblotting with anti-Flag antibodies (**a**) and anti-Myc antibodies (**b**). (**c**) DSS cross-linking assays of DJ-1 in HEK293T cells with Flag-tagged and Myc-tagged DJ-1 overexpression. (**d**) The schematic of deletion mutants at C terminus of DJ-1, from DJ-1 ΔC29 to DJ-1 ΔC6. (**e**) DSS cross-linking assays of DJ-1 WT and deletion mutants at C terminus (ΔC29, ΔC23, ΔC17, ΔC15 and ΔC6) in HEK293T cells. (**f**) The schematic of deletion mutants at C terminus of DJ-1, from DJ-1 ΔC6 to DJ-1 ΔC1. (**g)** DSS cross-linking assays of DJ-1 WT and deletion mutants at C terminus (ΔC6, ΔC5, ΔC4, ΔC3, ΔC2 and ΔC1) in HEK293T cells.

It is reported that the C-terminal G-helix-kink-H-helix motif is essential for protein stability and survival promoting activity of DJ-1 [26]. We explored how the C-terminal region had effects on the homodimerization of DJ-1. Different deletion mutations including DJ-1 ΔC6, ΔC15, ΔC17, ΔC23, ΔC29 in its C terminus were established according to the G-helix-kink-H-helix motif (Fig. 1d). The dimeric DJ-1 was completely disturbed in the ΔC29 mutant, which was consistent with previous report [26]. More interestingly, the DSS-mediated cross-linking assays showed the same results as DJ-1 ΔC29 mutant when we narrowed down the deletion region to ΔC23, ΔC17, ΔC15 and even to ΔC6 mutants, indicating that the last six amino acids at the C terminus were important to DJ-1 homodimerization (Fig. 1e). Next, we established the deletion mutants one by one among the last six amino acids, including DJ-1 ΔC1-ΔC6 (Fig. 1f) and it presented that once deleting more than two amino acids, the homodimerization of DJ-1 would be affected seriously because DJ-1 ΔC3 had no dimeric formation. When deleting two amino acids K188 and D189, the homodimerization was partially influenced, while there was no impact when D189 residue was deleted (Fig. 1g). Thus, our data suggest that motifs in C terminus, especially the L187, K188 and D189 residues regulate dimeric DJ-1 formation.

### Hydrophobic L187 residue is critical to DJ-1 homodimerization

The C terminus of DJ-1 is consist of many hydrophobic amino acids which may be critical for the homodimerization of DJ-1. To further study which residue among L187, K188 and D189 plays the most important role in the dimeric DJ-1 formation, we established different point mutants based on the three residues according to their property of charge (Fig. 2a). Given that the endogenous DJ-1 protein might have effects on the homodimerization of ectopic DJ-1, we utilized the DJ-1^-/-^ mouse embryonic fibroblast cells (MEFs) to exclude the interference of endogenous DJ-1. When we reintroduced human DJ-1 WT and its point mutants into DJ-1^-/-^ MEFs, we found that D189A and D189L mutants had little impact on DJ-1 homodimerization as their dimeric DJ-1 level was similar to wild type DJ-1 (Fig. 2b). And the homodimerization of K188E and K188R DJ-1 mutants was partially decreased according to the DSS-mediated cross-linking assay (Fig. 2b). Interestingly, when we reintroduced L187E mutant into DJ-1^-/-^ MEFs, the dimeric DJ-1 totally disappeared, which was the same as DJ-1 ΔC3 (Fig. 2b). The V51 residue is important to the homodimerization based on the crystal structure, which serves as a bridge between two monomeric DJ-1 [11, 24–26]. Then we introduced the V51C mutant as a control to further study the importance of C terminus to DJ-1. When DJ-1 ΔC3 and L187E mutants were reintroduced along with WT and V51C mutant into DJ-1^-/-^ MEFs, we found that V51C mutant completely interfered with homodimerization of DJ-1 as expected (Fig. 2c). And DJ-1 ΔC3 and L187E mutants also absolutely suppressed the dimeric DJ-1 formation like V51C mutant (Fig. 2c).

**Fig. 2.**
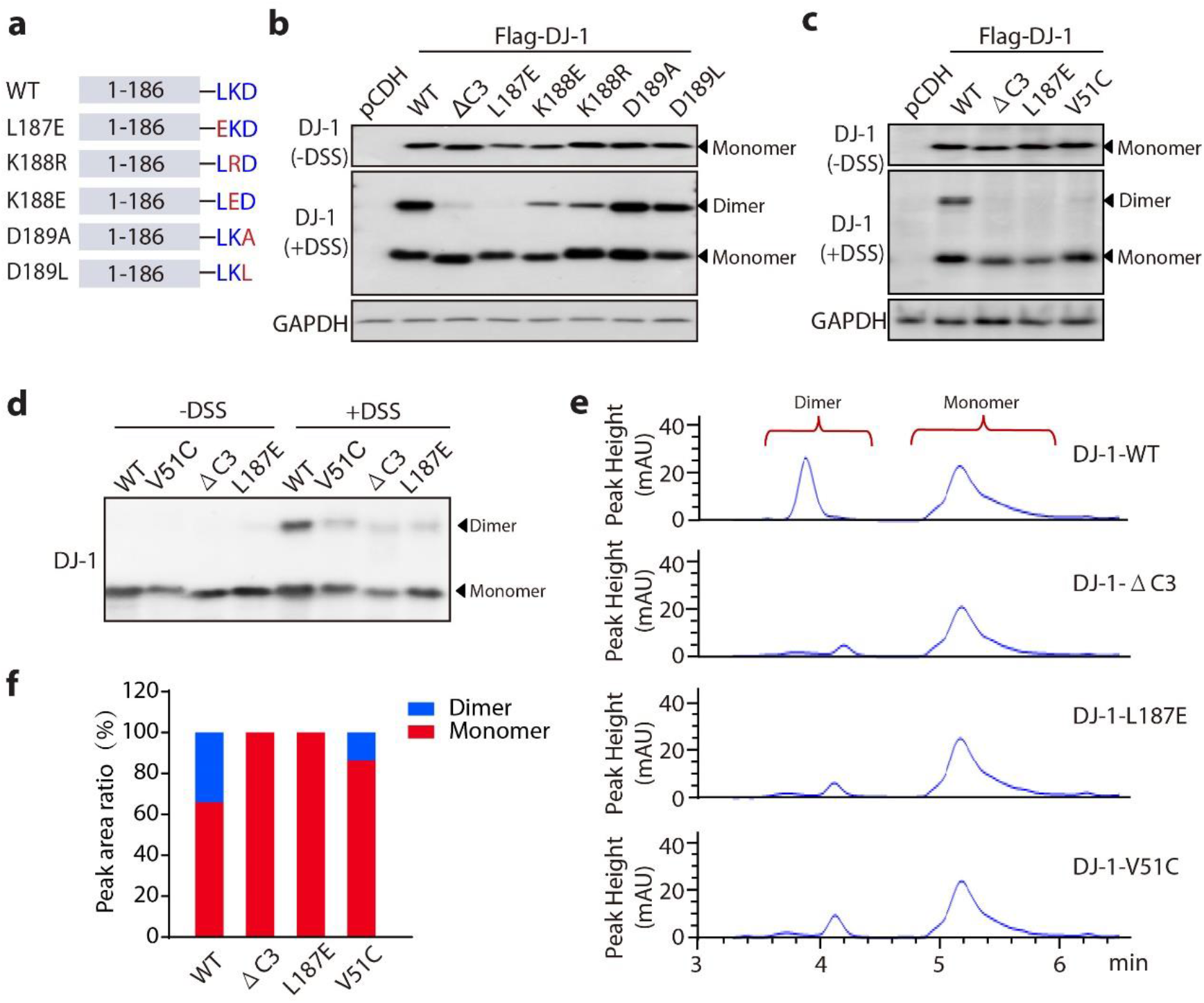
Hydrophobic L187 residue is critical to DJ-1 homodimerization. (**a**) The schematic of point mutants at C terminus of DJ-1. (**b**) DSS cross-linking assays of DJ-1 WT and point mutants at C terminus in DJ-1^-/-^ MEFs with re-introducing DJ-1 proteins. (**c**) DSS cross-linking assays of DJ-1 WT and mutants (DJ-1 ΔC3, L187E, V51C) in DJ-1^-/-^ MEFs with re-introducing DJ-1 proteins. (**d**) DSS cross-linking assays of purified DJ-1 WT and mutants (DJ-1 ΔC3, L187E, V51C) proteins. (**e**) Size exclusion chromatography analysis of purified DJ-1 WT and mutants (DJ-1 ΔC3, L187E, V51C) proteins. (**f**) Quantitative analysis of (**e**).

After investigating the influence of C terminus on the DJ-1 homodimerization at the cellular level, we asked whether purified protein had the same phenomenon. We cloned wild type DJ-1 and its ΔC3, L187E and V51C mutants into pET28a vectors, which express N-term 6×His-tagged. Then the cloned vectors were transformed into E. coli *BL21* (DE3) to express different DJ-1 proteins. The expressed DJ-1 proteins were purified by Ni2^+^-NTA column and the eluent was detected by SDS-PAGE (Fig. S1a–S1d). The eluent containing DJ-1 proteins were then subjected to dialysis in phosphate-buffered saline and the purity was identified. The coomassic brilliant blue stain result showed that the proteins were purified adequately (Fig. S1e). Next, we utilized the purified proteins to take DSS-mediated cross-linking assay, the result indicated that three mutants of DJ-1 inhibited the dimeric formation significantly compared with DJ-1 WT (Fig. 2d). Besides, the result of size exclusion chromatography showed that about 33% of protein was dimer in DJ-1 WT protein while the ratio in DJ-1 V51C mutant was nearly 15 %, and there was few dimeric DJ-1 detected in DJ-1 ΔC3 and L187E mutants when the detected smallest peak area was set as 50 mAU*s. (Fig. 2e and 2f). These *in vitro* results, together with the data in cell levels, further confirm that the C terminus of DJ-1, especially hydrophobic L187 residue is critical to homodimerization of DJ-1.

### C terminus of DJ-1 affects its deglycation activity

MGO produced by glucose oxidation, lipid peroxidation and DNA oxidation are reactive metabolites that form adducts on cysteine, lysine and arginine residues of protein [30]. MGO is converted into D-lactate by two enzymes Glyoxalase1 (Glo1) and Glyoxalase 2 (Glo 2) in the glyoxalase system via a mechanism requiring reduced glutathione (GSH) [31]. DJ-1 have been characterized as another glyoxalase in vitro, directly converting MGO into lactate without GSH (Fig. 3a) [18]. Later, Richarm group suggested that this glyoxalase activity is actually the deglycase activity, enzymatically removing MGO adducts from protein side-chains such as cysteine (Fig. 3b) [22]. The glyoxalase and deglycase activity could be summarized as the MGO detoxification ability of DJ-1 [30]. What we next wanted to investigate was that whether C terminus of DJ-1 has effects on the MGO detoxification ability of DJ-1. Firstly, we performed the glyoxalase assay utilizing purified protein and the result showed that the consumption of MGO presented an enormous difference between wild type DJ-1 and its mutants. The MGO consumption was significantly decreased in purified DJ-1 mutants compared with purified wild type DJ-1, suggesting the glyoxalase activity was dramatically inhibited in both DJ-1 ΔC3 and L187E mutants, and the suppression effect was demonstrated the same as V51C mutant (Fig. 3c). Then we investigated the deglycase activity of DJ-1 ΔC3 and L187E mutants. The absorbance at 288 nm represented the level of hemithioacetal, the intermediate in the reaction of MGO and N-acetylcyseine, which could be ultimately converted into lactate by DJ-1. The result showed that the decrease of hemithioacetal was obviously inhibited in DJ-1 ΔC3, L187E and V51C mutants, indicating that the deglycase activity of these mutants was suppressed (Fig. 3d). These data implied that the MGO detoxification ability of DJ-1 was regulated by its C terminus and hydrophobic L187 residue in vitro.

**Fig. 3.**
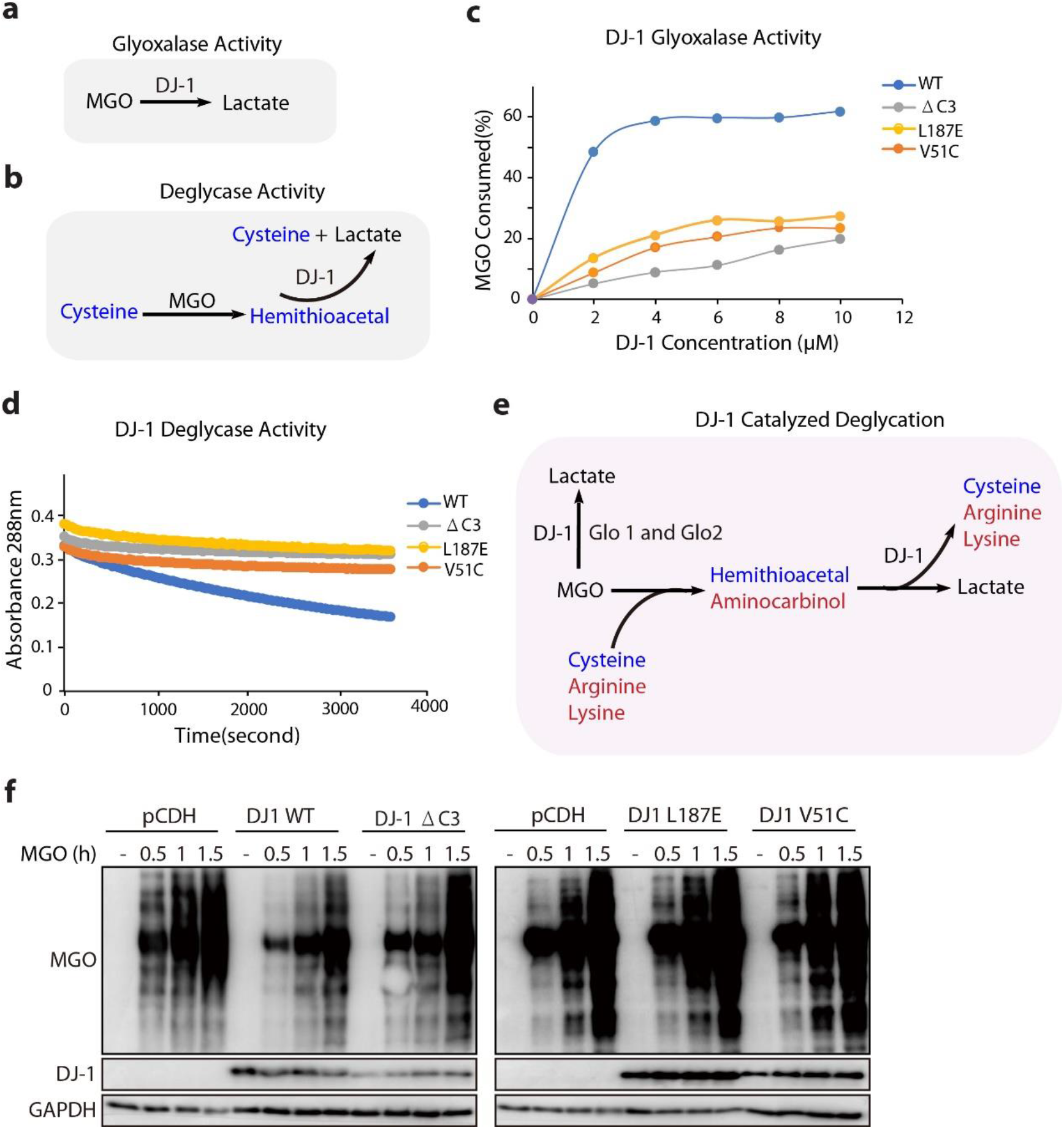
C terminus of DJ-1 affects its deglycation activity. (**a-b**) The schematic of DJ-1 glyoxalase activity (**a**) and deglycase activity (**b**). (**c**) Glyoxalase assay of purified DJ-1 WT and mutants (DJ-1 ΔC3, L187E, V51C), absorbance (540 nm for MGO) was measured. (**d**) Deglycase assay of purified DJ-1 WT and mutants (DJ-1 ΔC3, L187E, V51C), absorbance (288 nm for hemithioacetal) was measured for 60 min. (**e**) The schematic of DJ-1 catalyzed deglycation in DJ-1^-/-^ MEFs with re-introducing DJ-1 proteins. (**f**) DJ-1^-/-^ MEFs with reintroducing different human DJ-1 proteins were treated with 2Mm MGO for 0.5, 1 and 1.5 h, cells were harvested and the expression of DJ-1 and MGO-glycated protein were assayed by western blot. The relative gene expression is normalized to GAPDH.

Next, we studied the influence of DJ-1 C terminus on the MGO detoxification ability in cells. Besides over-expressed DJ-1 protein, the Glo1 and Glo2 in the glyoxalase system also convert MGO into lactate [31]. And MGO-glycated proteins containing cysteine, arginine and lysine residues could be only repaired by DJ-1 through different intermediate metabolites (Fig. 3e). We thus established stable over-expressed WT and a series of mutations DJ-1 in DJ-1^-/-^ MEFs to test the influence of DJ-1 C terminus on the MGO detoxification ability. After treated with 2 mM MGO, the cells were centrifuged and lysed for detecting the level of MGO-glycated proteins by a specific anti-MGO antibody. As expected, our result suggested that MGO was consumed distinctly by wild type DJ-1, indicating that DJ-1 WT has MGO detoxification ability in cells (Fig. 3f). Of note, the MGO level was partially reversed in DJ-1 ΔC3 mutant and absolutely reversed in DJ-1 L187E mutant (Fig. 3f), suggesting C terminus of DJ-1 affects its MGO detoxification ability in cells.

### The C terminus of DJ-1 regulates its suppression of ferroptosis

Given that recent study has suggested that DJ-1 could suppress ferroptosis triggered by lipid ROS, the impact of DJ-1 C terminus on the suppression of ferroptosis was next investigated [14]. We utilized Erastin, a classic ferroptosis inducer, to build the model of ferroptosis. DJ-1 WT and three mutants including DJ-1 ΔC3, L187E and V51C were reintroduced into DJ-1^-/-^ MEFs (Fig. 4a). After treated with Erastin for 12 h, the cells were centrifuged to perform flow cytometer using the fluorescent probe C11-BODIPY. Over-expression of wild type DJ-1 significantly inhibited Erastin-induced accumulation of lipid ROS, a classic biomarker of ferroptosis, and V51C mutant had partially inhibition effects. While DJ-1 ΔC3 and L187E mutants had fairly effects on the accumulation of lipid ROS induced by Erastin (Fig. 4b and 4c).

**Fig. 4.**
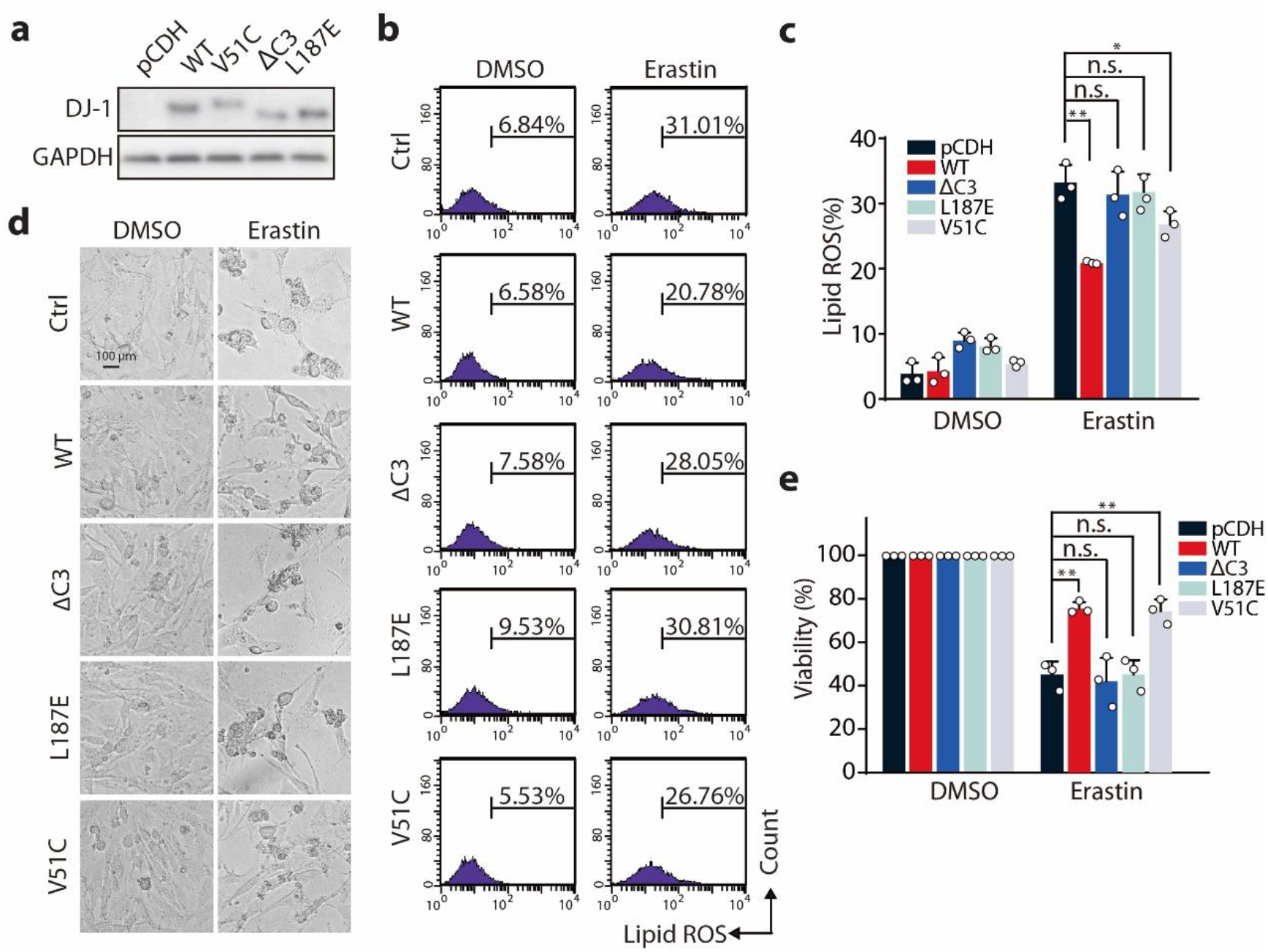
The C terminus of DJ-1 regulates its suppression of ferroptosis. (**a**) Western blot analysis of DJ-1 expression in DJ-1^-/-^ MEFs with re-introducing human DJ-1 WT and mutation proteins. (**b**) Indicated MEFs were treated with Erastin (400 nM) for 12 h, and lipid ROS production was assayed by flow cytometry using C11-BODIPY and representative data are shown. (**c**) Quantitative analysis of (**b**) and the statistical analysis is from three independent experiments. (**d**) Phase-contrast images of indicated MEFs treated with 400 nM Erastin for 24 h. Independent experiments were repeated three times and representative data are shown. (**e**) Cell viability was assayed in indicated MEFs treated with 400 nM Erastin for 24 h. All data are representative of three independent experiments and the error bar indicates the s.d. value. *: p < 0.05; **: p < 0.01.

Moreover, the effect of DJ-1 on Erastin-triggered ferroptotic cell death was also examined. As shown in Figure 4d and 4e, wild type DJ-1 displayed an inhibitory role in the effect of Erastin triggered ferroptotic cell death, and the suppression of ferroptosis was eliminated when DJ-1 had mutations DJ-1 ΔC3, L187E at the C terminus. And DJ-1 V51C mutant remained part of the suppression of ferroptosis, which was consistent with the result of lipid ROS level (Fig. 4d and 4e). Therefore, our results further implied that C terminus of DJ-1 regulates its suppression of ferroptosis and when its C-terminal structure was destroyed, its inhibition of ferroptosis would be abolished.

## Discussion

DJ-1 is a multi-functional protein associated with both neurodegeneration and neoplasia and it is believed that DJ-1 carries out its function exclusively in the dimeric state [32–34]. In our previous study, the dimeric DJ-1 depolymerizes among the apoptosis regulated by 4-HPR, indicating that the homodimerization of DJ-1 may be assuredly critical to its function [15]. The pathogenic L166P mutant of DJ-1 is responsible for Parkinson disease (PD) because it impairs the neuronal cytoprotective function of DJ-1 by jeopardizing dimer formation and protein stability [35]. It is interesting to investigate whether the loss of function is caused by disappearance of homodimerization or is due to the impacts on DJ-1 protein stability in these mutants. Our studies showed that monomeric DJ-1 ΔC3 and L187E mutants were stable because no significant protein degradation was observed compared with DJ-1 WT both in cells and *in vitro*, differing from L166P mutant. These results indicated that the level of homodimerization could not fully represent the stability of DJ-1 protein and the mutants DJ-1 ΔC3 and L187E are more useful tools to carry out studies on the function of dimeric DJ-1.

It is reported that G-helix and kink motif are important because they determine the C-terminal helix-kink-helix, which is essential for protein stability and survival promoting activity of DJ-1 [26]. In addition, the pathogenic L166P mutant induces a loss of DJ-1 function and instead of forming a dimeric structure, it may assemble in high molecular weight oligomers [35, 36]. As for the structural of DJ-1, L166 is located at the middle of G-helix motif and its mutation breaks the helix [35]. The hydrophobic interactions with a series of residues located at the subunit-subunit interface including V181, K182, and L187 of the C terminus may affect the stability of the homodimer [37]. However, this hypothesis has not yet been proven by the analysis of the conformational changes or the experiments in vitro. In this study, we confirmed that wild type DJ-1had homodimerization and the C terminus was crucial for the stability of dimeric DJ-1 protein. The critical motif of the C terminus was narrowed down among L187, K188, D189 and further research showed that hydrophobic L187 residue played the most important role in DJ-1 homodimerization. And these data supported the hypothesis that hydrophobic interactions may had effects on the stability of the DJ-1 homodimer, prompting that studies on other hydrophobic residues located at the subunit-subunit interface are worth carrying out. Moreover, probes designed based on the hydrophobic C terminus of DJ-1 are useful tools to carry out researches on DJ-1 structure and function. The inhibitors of dimeric DJ-1 which may have anti-tumor synergistic effect could be designed according to the structure of the hydrophobic C terminus.

As for the studies on the function of DJ-1, the C terminus was responsible for MGO detoxification and suppression of ferroptosis. The glyoxalase and deglycase activity repair MGO- and GO-glycated nucleotides and proteins, making DJ-1display a cytoprotective role [18, 21, 22, 31]. We found that the hydrophobic C terminus, especially L187 residue was crucial for DJ-1 MGO detoxification because both deglycase and glyoxalase were totally inhibited in L187E mutant *in vitro*. While the effects in cells seemed not to exactly accord with the results *in vitro* as DJ-1 ΔC3 had residual effects of the deglycation activity. We think that it was probably because the effects of DJ-1 was complicated for its multi-functions in cells. In addition, Glo 1 and Glo 2 took part in MGO detoxification and the level of Glo 1 and Glo 2 also impacted the MGO level. Given that the crucial influence of C terminus on the MGO detoxification, the *in vitro* deglycase activity and glyoxalase activity assay can be applied to screen the inhibitors of dimeric DJ-1 with the high efficiency.

In addition, the results about the suppression of ferroptosis were consist with MGO detoxification ability as DJ-1 ΔC3 and L187E mutants lost the inhibition of ferroptosis compared with DJ-1 WT. Besides, the suppression was more obvious in DJ-1 ΔC3 and L187E mutants than V51C mutant, indicating that the hydrophobic C terminus may have other effects such as involving regulation of signaling pathway [4, 6, 38]. All these data supported our previous discovery that DJ-1 was a negative modulator of ferroptosis providing new opportunities to facilitate ferroptosis-based cancer therapy [14].

Taken together, this study found that the hydrophobic C-terminal region of DJ-1 regulated its homodimerization, deglycation activity and suppression of ferroptosis. It is a valuable supplement to the existing studies about the relationship between the structure and functions of DJ-1. It also provides useful ideas and direction on designing and screening inhibitors of DJ-1, which may have potential in anti-tumor effects [39, 40].

## Acknowledgement

This work was supported by grants from National Science & Technology Major Project “Key New Drug Creation and Manufacturing Program”, China (2018ZX09711002 to Hong Zhu), the National Natural Science Foundation of China (No. 81773757 to Meidan Ying; No.81402951 to Ji Cao), and Zhejiang Medical and health science and technology program (No. 2020384517 to Chengyong Du).

## Author contributions

Ji Cao, Li Jiang, and Xiaobing Chen designed the research; Ji Cao, Li Jiang, and Xiaobing Chen wrote the manuscript; Li Jiang,Xiaobing Chen, Qian Wu, Haiying Zhu, Chengyong Du performed the biochemical and cellular studies; Ji Cao, Li Jiang, Xiaobing Chen, Hong Zhu, and Meidan Ying analyzed the results; Ji Cao, Hong Zhu, Meidan Ying, Qiaojun He and Bo Yang directed the study.

---

**Supplementary Fig. S1.**
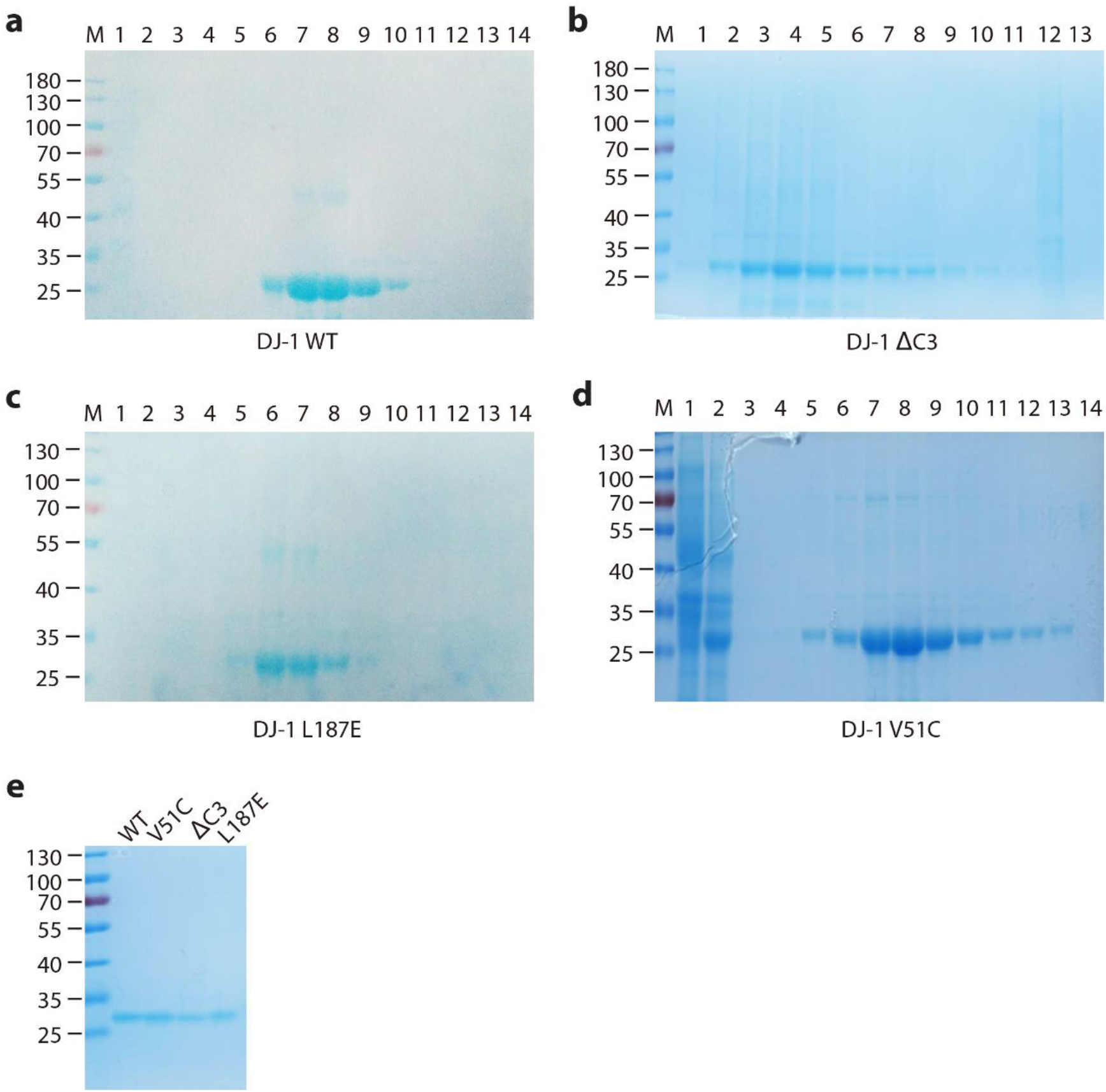
(**a**-**d**) Coomassie brilliant blue staining of the fractions from Ni^2+^-NTA column of DJ-1 proteins, including DJ-1 WT (**a**), DJ-1 ΔC3 (**b**), DJ-1 L187E (**c**) and DJ-1 V51C (**d**). (**e**) Coomassie brilliant blue staining of purified DJ-1 proteins after dialysis.

**Supplementary Table S1:**
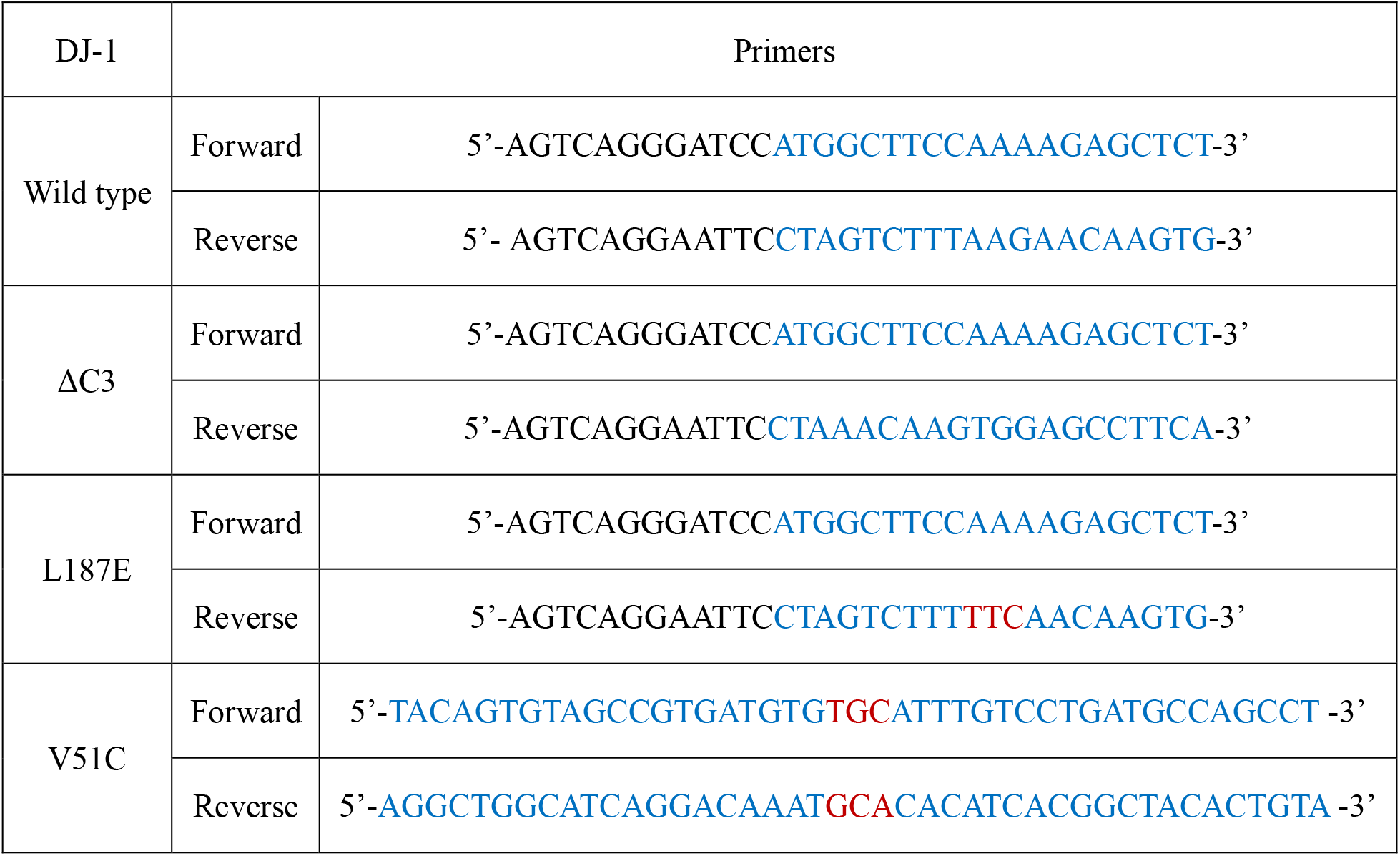
Sequences of wild type DJ-1 and mutagenesis primers with N-terminal 6×His-tag for pET28a vector.

**Supplementary Table S2:**
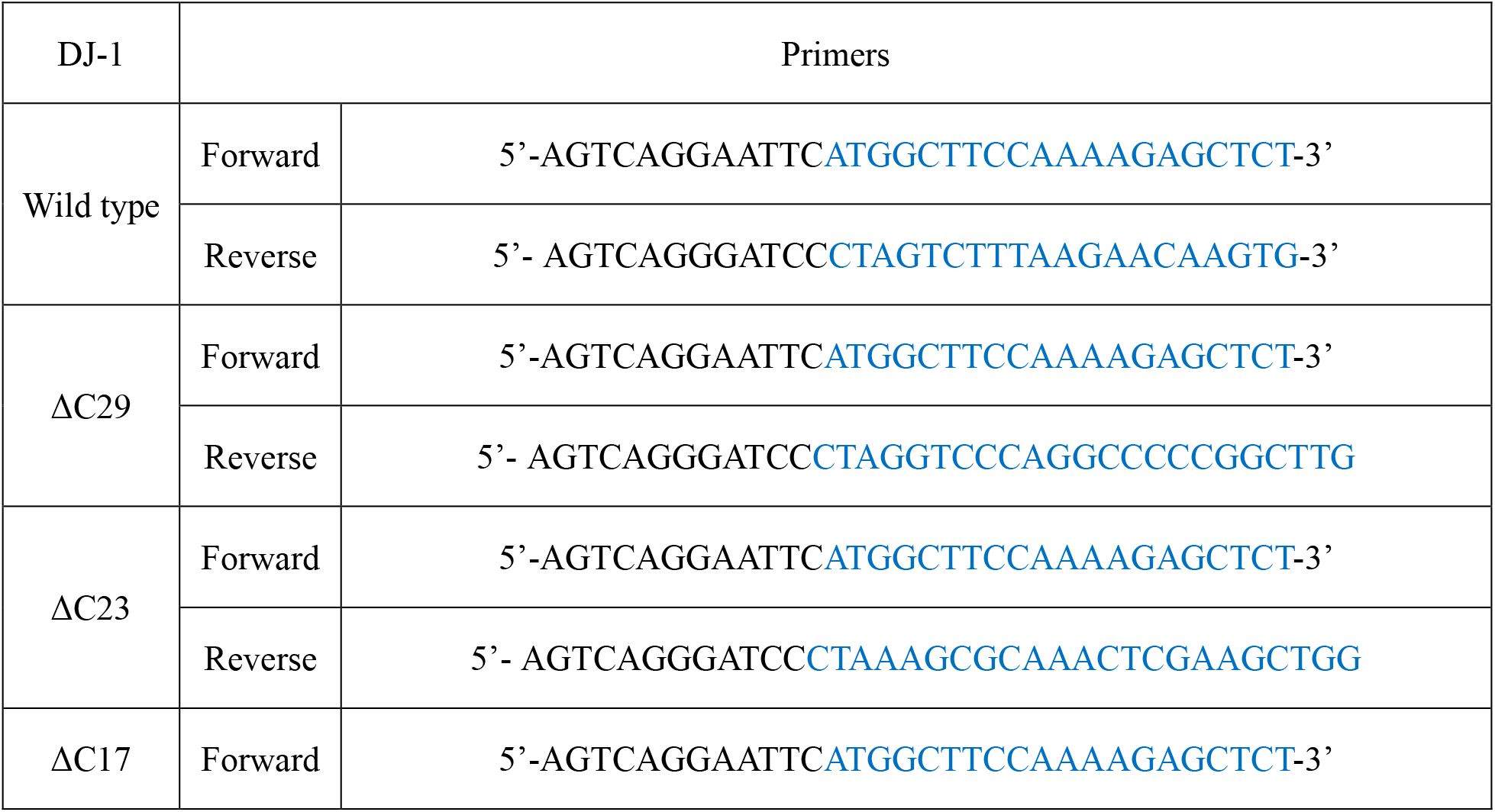
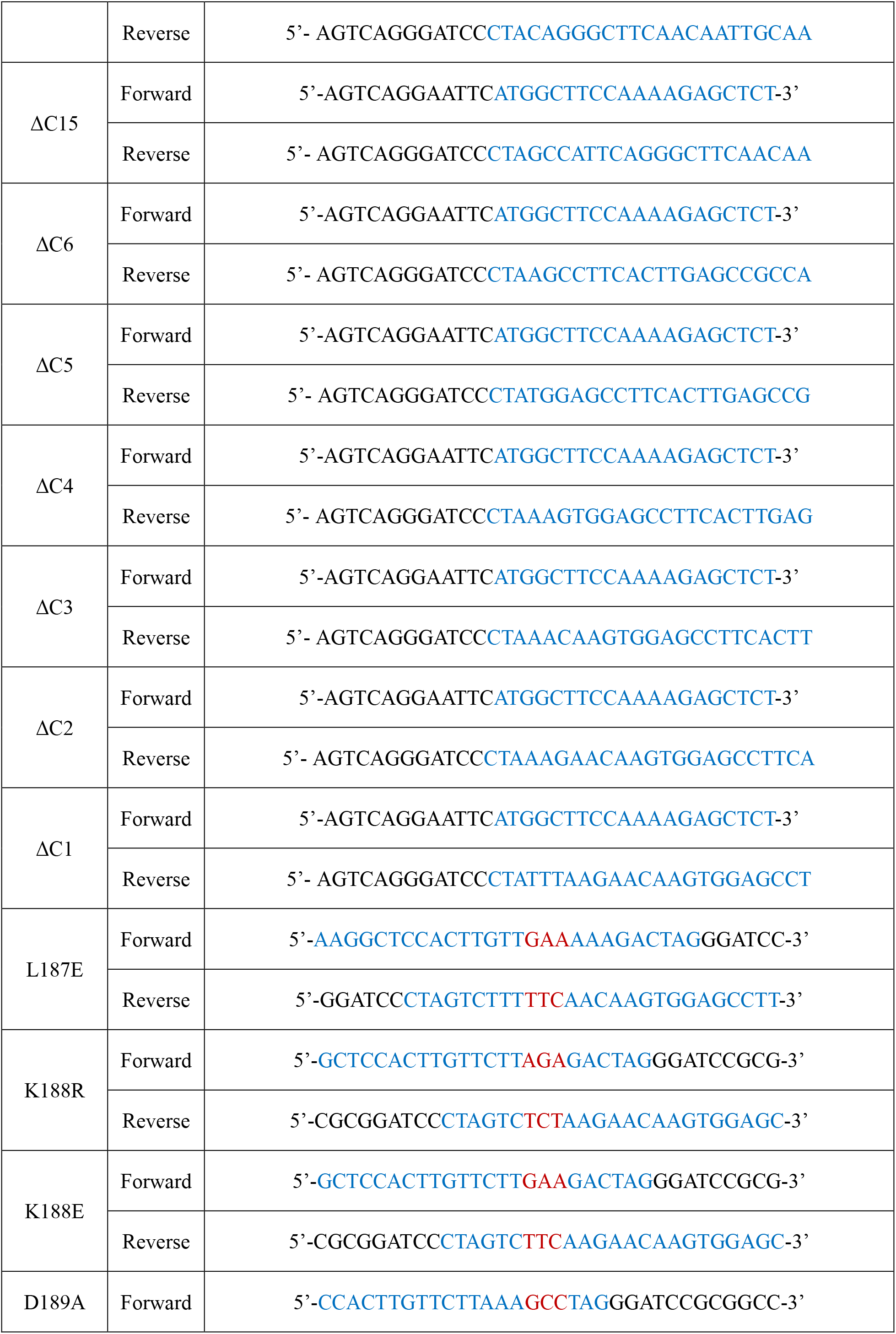
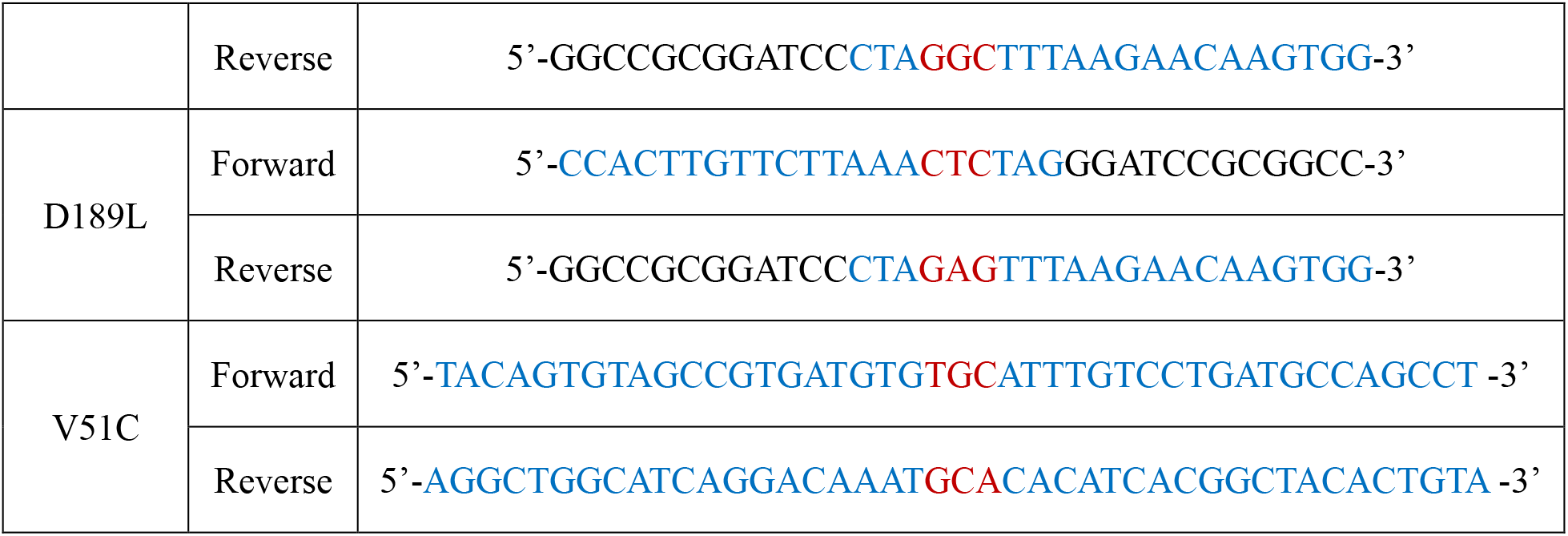
Sequences of wild type DJ-1 and mutagenesis primers with N-terminal Flag-tag for pCDH-EF1-Puro plasmid.

