## Supplementary figures and images for "C Terminus of DJ-1 Determines Its Homodimerization, Deglycation Activity and Suppression of Ferroptosis"

### Supplemental Table 1

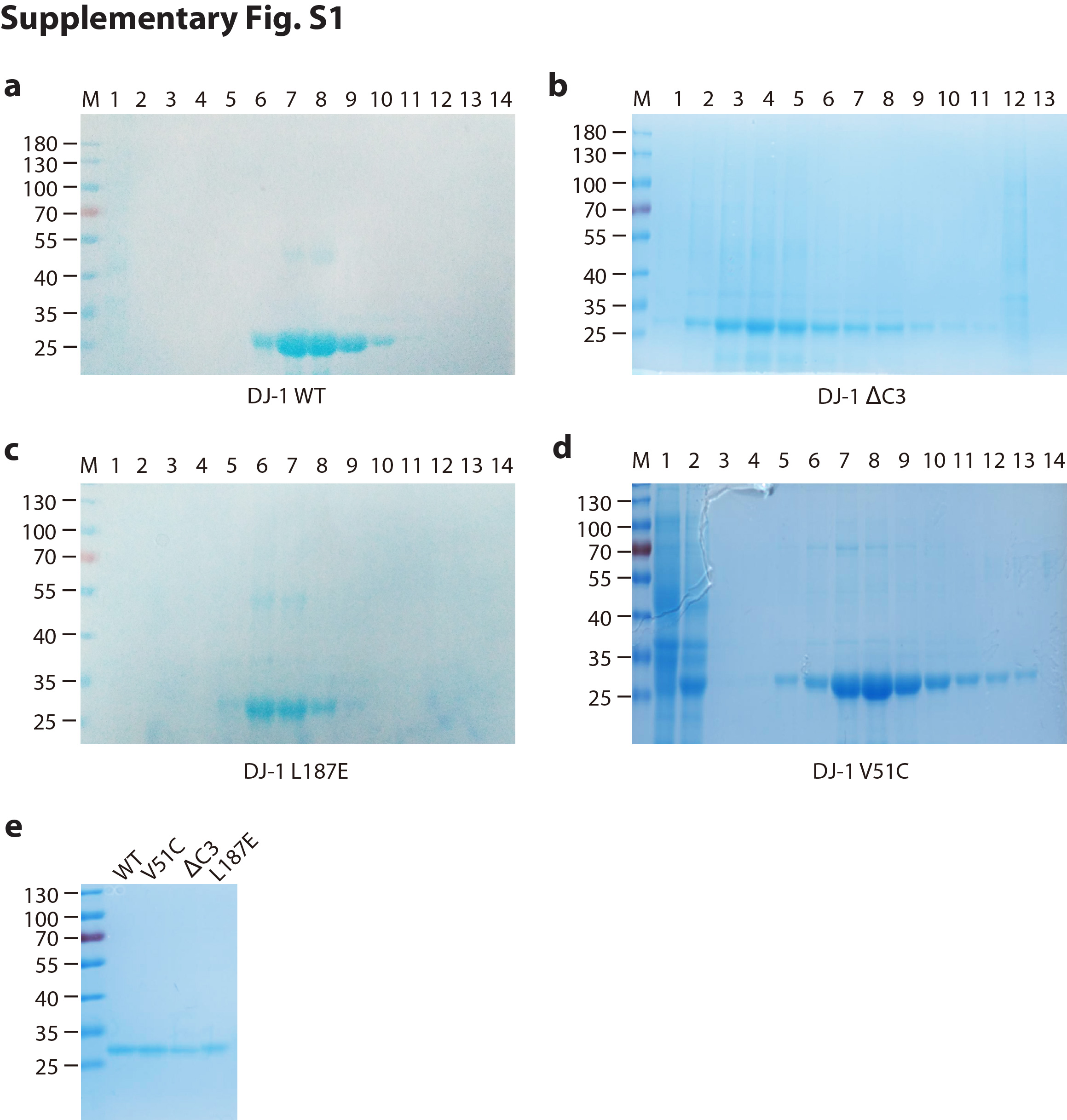
